# Improved neurocognitive performance in FIV infected cats following treatment with the p75 neurotrophin receptor ligand LM11A-31

**DOI:** 10.1101/2020.06.17.156596

**Authors:** Jonathan E. Fogle, Lola Hudson, Andrea Thomson, Barbara Sherman, Margaret Gruen, B. Duncan Lacelles, Brenda M Colby, Gillian Clary, Frank Longo, Rick B Meeker

## Abstract

HIV rapidly infects the central nervous system (CNS) and establishes a persistent viral reservoir within microglia, perivascular macrophages and astrocytes. Inefficient control of CNS viral replication by antiretroviral therapy results in chronic inflammation and progressive cognitive decline in up to 50% of infected individuals with no effective treatment options. Neurotrophin based therapies have excellent potential to stabilize and repair the nervous system. A novel non-peptide ligand, LM11A-31, that targets the p75 neurotrophin receptor (p75^NTR^) has been identified as a small bioavailable molecule capable of strong neuroprotection with minimal side effects. To evaluate the neuroprotective effects of LM11A-31 in a natural infection model, we treated cats chronically infected with feline immunodeficiency virus (FIV) with 13 mg/kg LM11A-31 twice daily over a period of 10 weeks and assessed effects on cognitive functions, open field behaviors, activity, sensory thresholds, plasma FIV, cerebrospinal fluid (CSF) FIV, peripheral blood mononuclear cell provirus, CD4 and CD8 cell counts and general physiology. Between 12 and 18 months post-inoculation, cats began to show signs of neural dysfunction in T maze testing and novel object recognition, which were prevented by LM11A-31 treatment. Anxiety-like behavior was reduced in the open field and no changes were seen in sensory thresholds. Systemic FIV titers were unaffected but treated cats exhibited a log drop in CSF FIV titers. No significant adverse effects were observed under all conditions. The data indicate that LM11A-31 is likely to be a potent adjunctive treatment for the control of neurodegeneration in HIV infected individuals.

**Author Summary:** There are no effective treatments to halt the progression of most neurodegenerative diseases including HIV-associated neurodegeneration. Neurotrophins have the potential to provide strong neuroprotection but it has been difficult to develop usable interventions. A new drug, LM11A-31, that targets the p75 neurotrophin receptor has been developed that provides potent neuroprotection, is orally bioavailable and has the potential to prevent disease progression. The current studies were designed to evaluate the effects of the compound in an animal model of active HIV infection in preparation for a human clinical trial. Treatment of chronically infected animals with LM11A-31 normalized deficits in T maze performance, novel object recognition and open field behavior with no measurable adverse effects. Potential adverse effects associated with natural neurotrophins such as changes in sensory perception and increased systemic viral burden were not observed. A decrease in CSF FIV titers and a slight improvement in the CD4:CD8 ratio suggested that LM11A-31 may have beneficial effects beyond the anticipated neuroprotective effects. These findings are similar to beneficial effects seen in other animal models of neurodegeneration and CNS injury and support the use of LM11A-31 as an adjunctive neuroprotective agent for the treatment of HIV infected individuals.

## Introduction

Lentiviruses such as HIV, SIV and FIV rapidly penetrate the central nervous system (CNS) where they establish perivascular macrophage reservoirs and infect microglia and astrocytes. Sustained virus production by microglia and macrophages is difficult to treat with antiretroviral drugs due to poor penetration of the blood brain barrier. The resulting inflammation leads to neural dysfunction and an accelerated aging phenotype(1–8) with varying degrees of cognitive decline in up to 50% of infected individuals(9–11). Cognitive decline adversely affects quality of life and there are currently few therapeutic options available. Attempts to protect the nervous system are focused, in part, on interventions that reduce damage due to HIV-associated inflammation. We and others have shown that activation of mononuclear phagocytes (microglia and macrophages) results in the release of factors that induce a gradual accumulation of intraneuronal calcium(12–19). In vitro models of neuroinflammation, show that chronic elevation of calcium is quickly followed by cytoskeletal damage, transport deficits and focal swelling (beading) within dendrites and axons(20). These dendritic and axonal swellings accumulate mitochondria, endoplasmic reticulum (ER), p75 neurotrophin receptors (p75^NTR^) and a unique oligomeric form of Tau(21), as well as other proteins, providing an environment poised for mitochondrial dysfunction, ER stress and pathological protein modifications. Interestingly, an almost identical progression is seen in animal models of Alzheimer disease (AD) indicating that this is a general response to inflammation which renders neurons vulnerable to neurodegeneration(17, 22). Importantly, these changes are reversible with early intervention(20).

A number of in vitro studies have shown that neurotrophin signaling has the capacity to reduce calcium-induced changes and prevent cytoskeletal disruption (reduced beading, sparing of dendrites) that underlie the initial development of neuronal vulnerability in response to gp120 and other HIV/FIV virion components. For example, in vitro studies have clearly demonstrated that cell culture medium from macrophages or microglia treated with neurotrophins abrogates lentivirus-induced neurodegenerative changes(20, 23–28). To capitalize on the protective potential of neurotrophin signaling, the nerve growth factor (NGF) loop 1 mimetic, LM11A-31, was developed to provide a stable, small molecule ligand targeted to p75^NTR^(29, 30). This non-peptide ligand crosses the blood brain barrier and promotes neuroprotective signaling through the activation of pro-survival pathways and inhibition of degenerative pathways(31, 32). In vitro, LM11A-31 has neurotrophic and neuroprotective effects in neural cultures treated with HIV gp120(20) as well as models of Alzheimer disease(31, 32) and p75^NTR^-mediated apoptosis(33). A number of in vivo studies in animal models have clearly demonstrated that LM11A-31 prevents neural damage and cognitive decline in models of aging, Alzheimer disease, Huntington disease, traumatic brain injury and spinal cord injury with no reported adverse effects(32, 34–37). Thus, LM11A-31 has properties that strongly suggest it is useful as a disease modifying intervention to prevent CNS dysfunction and cognitive decline in HIV infected patients.

To assess LM11A-31 efficacy in an animal model of HIV, we treated FIV infected cats with LM11A-31 for a period of 10 weeks and assessed cognitive function, virus production, T cell numbers, sensory function, and general health relative to pretreatment and placebo controls. Untreated FIV infected cats have neuropathological changes similar to HIV infected humans although the changes are generally less severe(38) resembling the gradual progression in individuals on antiretroviral therapy. A subtle gliosis can be seen as early as 1 week post-inoculation followed later during late asymptomatic or pre-AIDS stages of disease by: diffuse gliosis, microglial nodules, meningitis, perivascular infiltrates, white matter pallor, multinucleated cells (listed in order from more common to more rare findings). Some neuron loss can be detected in asymptomatic FIV-infected cats at ∼3 years post infection and matches the pattern of loss seen in humans(39). Other changes include neurological dysfunction(40–44), cortical atrophy revealed by MRI(43), and discrete white matter lesions(41, 43). These changes indicate that the FIV model is a good model to examine effects of compounds on the gradual development of neurological disease in response to lentiviral infection. We therefore treated FIV infected cats with LM11A-31 at a time when cognitive deficits first begin to appear to determine if the compound could suppress disease progression.

## Results

### LM11A-31 achieves high CSF to plasma ratios and exhibits rapid clearance following intravenous or oral administration

To determine the distribution and half-life of the drug and screen for possible adverse effects, the drug was administered orally (n=2), intravenously (n=5) or subcutaneously in pluronic gel (n=5). LM11A-31 reached therapeutic concentrations in CSF for all dosing strategies. Drug delivered subcutaneously was retained longer but achieved lower CSF penetration with a CSF/plasma ratio of 0.257. Intravenous and oral delivery showed higher CSF penetration (average CSF:plasma ratio of 7.0 for both). Pharmacokinetic (pK) data following oral or intravenous delivery of 10 mg/kg LM11A-31 were similar (Fig. 1). Clearance was rapid in both plasma and CSF with t_1/2_ estimates of 1.3-2.1 hrs. The CSF concentration was maintained above the minimum target therapeutic concentration of 10 nM for approximately 3.7 hrs. Peak drug concentration and T_1/2_ estimates were similar to values obtained in mouse studies with a comparable dose, based on body surface area(32). No significant adverse effects were noted in response to the drug, although a transient decrease in heart rate was noted in some cats. Given these results and to match the proposed route of delivery in humans, we elected to dose the cats orally.

**Fig 1.**
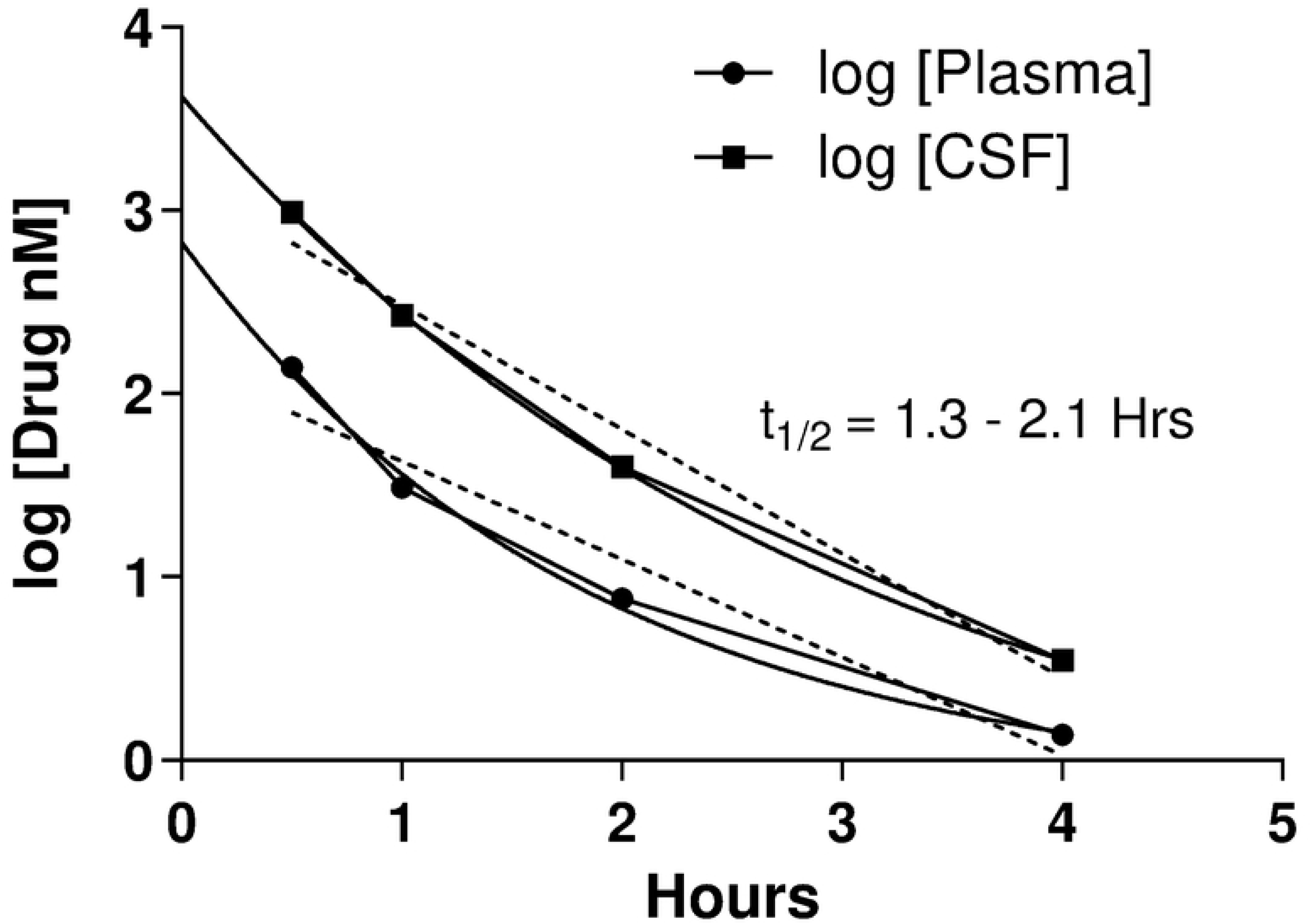
Pharmacokinetics of LM11A-31 in feline plasma and cerebrospinal fluid (CSF). An intravenous bolus of 10 mg/kg LM11A-31 was delivered at time 0 and the concentration of drug was measured in samples collected at 0.5, 1.0, 2.0 and 4 hours. The compound rapidly reached peak concentrations of 140-977 nM with higher concentrations in CSF than plasma. A similar rate of decline was seen in both compartments with an estimated t_1/2_ of 1.3-2.1 hours.

### Effect of LM11A-31 on plasma and CSF viral loads

Plasma FIV copies following intracranial inoculation were assessed for each cat (Fig. 2A). Samples collected just prior to inoculation were all negative for FIV. Two weeks after intracerebroventricular (icv) delivery of FIV, plasma FIV RNA rose to an average of log 5.86 ± 0.21 copies/ml in the group that subsequently received the placebo and log 5.18 ± 0.32 copies/ml in the group randomly selected to receive the drug. We observed a gradual decline in FIV viral loads over the next 18 months in both groups with no differences between groups, which is consistent with viral kinetics noted in our previous studies(45). Final FIV titers were measured after the completion of testing at approximately 20 months, while the cats were still being administered LMA11A-31. When FIV titers at the end of the 10 week treatment were compared to pretreatment levels, we found that plasma FIV rose slightly in the placebo treated cats and remained stable in the LM11A-31 treated cats; however, the changes were not significant. Cerebrospinal fluid (CSF) FIV followed a similar pretreatment pattern (Fig. 2B) with peak levels at 5.80 ± 0.10 log copies/ml and 4.82 ± 0.10 log copies/ml for the placebo and LM11A-31 groups respectively and a steady decline over time to similar pretreatment values of 2.63 (FIV) and 2.86 (FIV+LM11A-31) log copies/ml. Following treatment with LM11A-31, CSF FIV dropped significantly by 1.27 logs relative to the placebo group (p=0.037, Fig. 2B) suggesting that the drug may also reduce viral synthesis or perhaps facilitate FIV clearance from the CSF. FIV PBMC provirus declined over time with almost identical values of 3.64-3.39 log copies/10^6^ PBMCs at 18 months. Quantification of FIV DNA in the PBMCs during treatment with LM11A-31 (Fig. 2C) indicated there were no differences in the proviral burden before and after treatment. These findings indicate that treatment with a p75^NTR^ ligand had no significant effects on systemic viral titers but may have a small suppressive effect on virus titers in the CNS.

**Fig 2.**
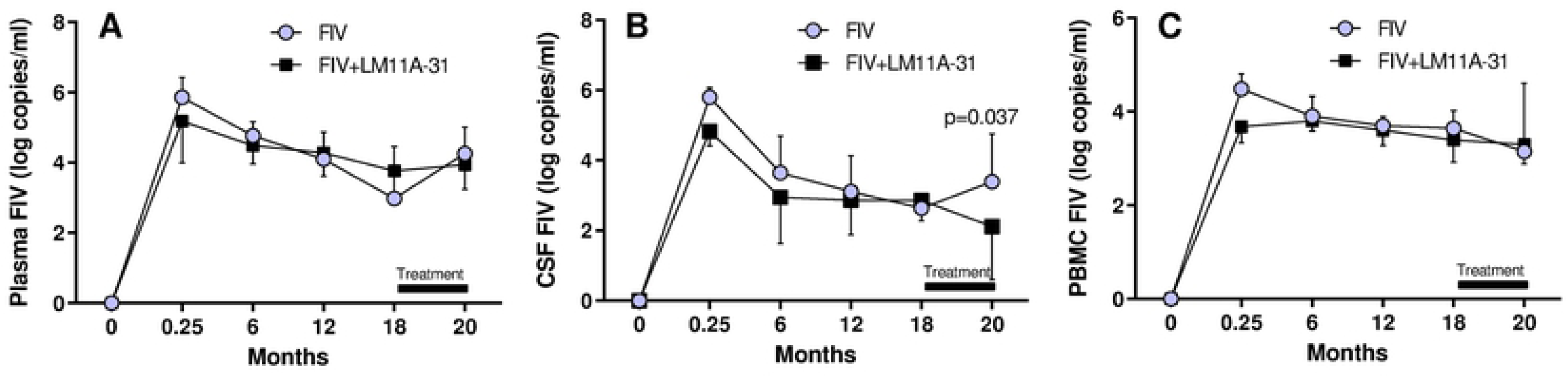
FIV RNA in plasma and CSF and FIV DNA in peripheral blood mononuclear cells (PBMCs) over the 20 months. A. FIV RNA in plasma peaked at 5.2-5.8 log copies/ml and decreased to 2.98-3.77 log copies/ml by 18 months with no significant differences between groups. B. FIV RNA in CSF peaked at 4.8-5.8 log copies/ml and dropped to 2.6-2.8 log copies/ml by 18 months. A significant decrease to 2.1 log copies/ml in the treated group (FIV+LM11A-31) was seen relative to an increase to 3.4 log copies/ml in the vehicle group (FIV) (unpaired T-test of pre to post differential, treated vs. untreated, p=0.032). C. FIV DNA in PBMCs peaked at 3.7-4.5 log copies/ml and thereafter remained relatively stable at 3.4-3.9 log copies/10^6^ PBMCs. No changes were seen after treatment with LM11A-31.

### Body Weights and body conditions cores were unaffected by LM11A-31 treatment

Body weight and body condition scores were monitored throughout the study to assess any adverse effects of infection or treatment. The data shown in Figure 3, illustrate that, with the exception of the expected acute drop in body weight immediately following infection (7-8%) and again at approximately 29-33 weeks (3-4%), all cats showed an overall body weight gain at the same rate. Treatment with LM11A-31 had no effect on body weight. Body condition scores paralleled the changes in body weight, again with no effect of treatment indicating that the cats were healthy and eating well throughout the experiment.

**Figure 3.**
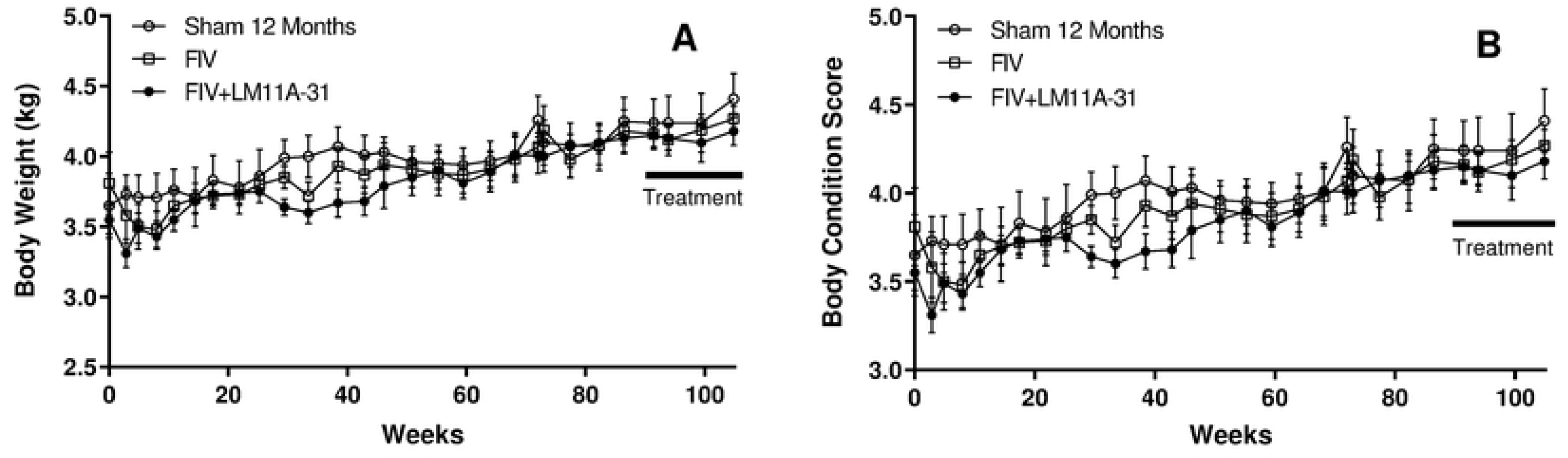
Average body weight and body condition scores. A. A small decrease in body weight was seen following infection and again for the infected cats at 29-33 weeks. By 38 weeks, rate of gain was restored and by 50 weeks body weights were the same for all groups. Treatment with LM11A-31 (bar) had no effect on body weight. B. Body condition scores paralleled the body weight data, again with no effect of treatment.

### The CD4+:CD8+ ratio improved after treatment

As expected, a decline in the CD4^+^:CD8^+^ T cell ratio was seen over time (Fig. 4A). The drop was similar in both FIV infected groups and was significant relative to sham inoculated controls. Treatment with LM11A-31 (20 month data) caused a small, but significant increase in the CD4:CD8 ratio (Fig. 4B) whereas both the placebo and sham groups showed small decreases, suggesting that LM11A-31 may have a favorable effect upon T cell population dynamics. These changes were too small to interpret the physiological relevance, but additional studies should determine if this trend continues with a longer treatment duration.

**Fig 4.**
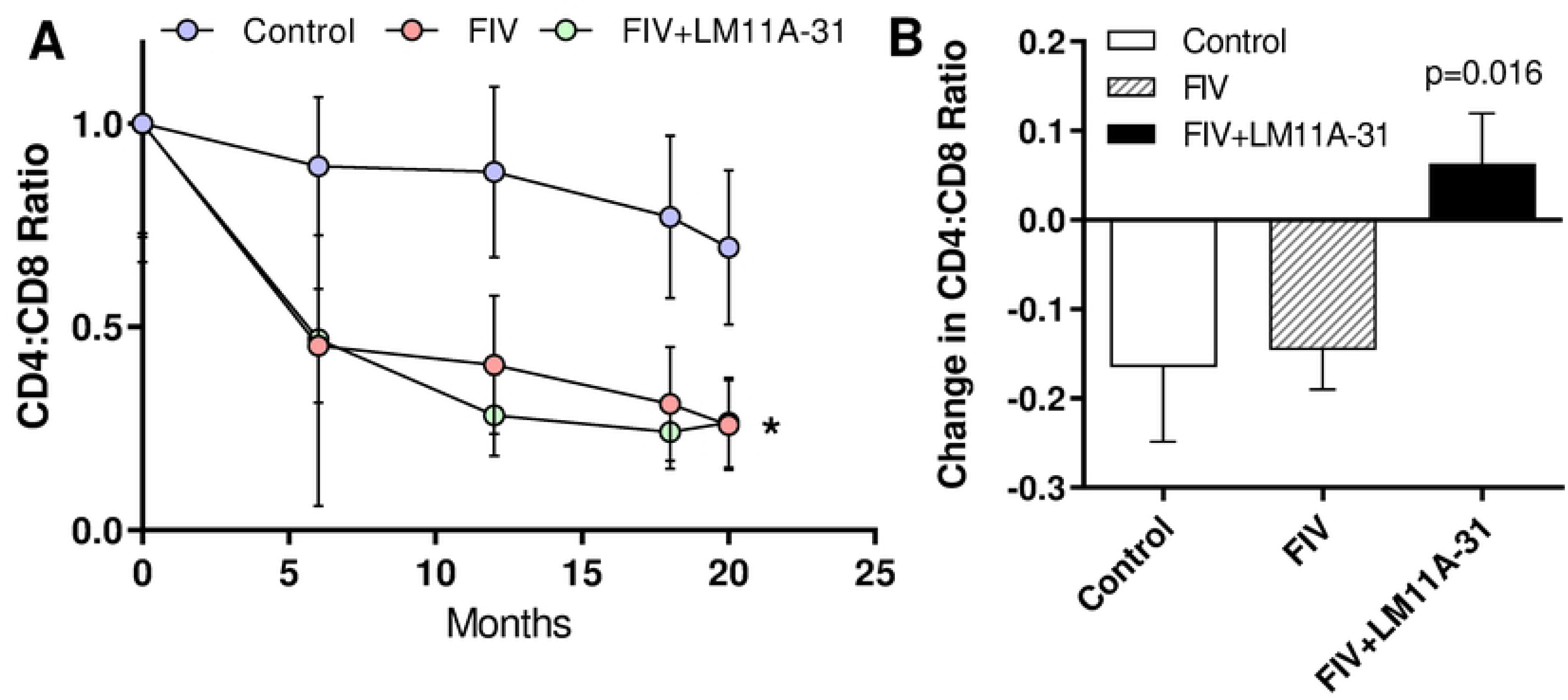
Ratio of CD4:CD8 T cells over time. A. As expected, we observed a large drop in the CD4:CD8 ratio in the FIV infected cats relative to sham inoculated controls. B. After treatment with LM11A-31, we noted a small increase in the CD4:CD8 ratio relative to placebo treated cats (*unpaired t-test of changes in placebo vs. treated cats, p=0.016).

### No adverse effects were observed in response to drug administration or FIV infection

Before initiating the behavior studies reported here, safety studies were performed in a subset of cats. Cats were administered the drug orally (n=4, LM11A -31 10 mg free base/kg) and assessed before and 48 hours after dosing; intravenously (n=2, LM11A 10 mg free base/kg) and assessed before and 24 hours after dosing; and subcutaneously (n=2, LM11A 10mg free base/kg) and assessed before and 72 hours after dosing. In these safety studies, a decreased hematocrit was noted in 2 of 4 cats at 48 hours following oral dosing, in 1 of 2 cats at 72 hours following intravenous dosing and in 2 of 2 cats following subcutaneous dosing. However, the effect was transient. The cats in the current longitudinal study reported here exhibited only sporadic, mild decreases in hematocrit over the entire study period (both control and treated groups). Further, no other significant laboratory changes were noted on the CBC, serum biochemistry profile or urinalysis throughout the entire length of the study.

### Daily activity was unaffected by treatment with LM11A-31

A subset of sham inoculated (n=5) and FIV infected (n=6) were monitored for changes in activity using Actical monitors over a period of 24 days. No other testing was administered during this period. On average, FIV infected cats at 18 months post-inoculation showed less total daily activity than matched sham controls (Fig. 5). LM11A-31 treatment over a 17-day period had no effect on average daily activity. Activity patterns were similar across cats with regard to time. The pattern of activity did not change significantly after introduction of LM11A-31 in the sham controls or FIV infected cats.

**Fig 5.**
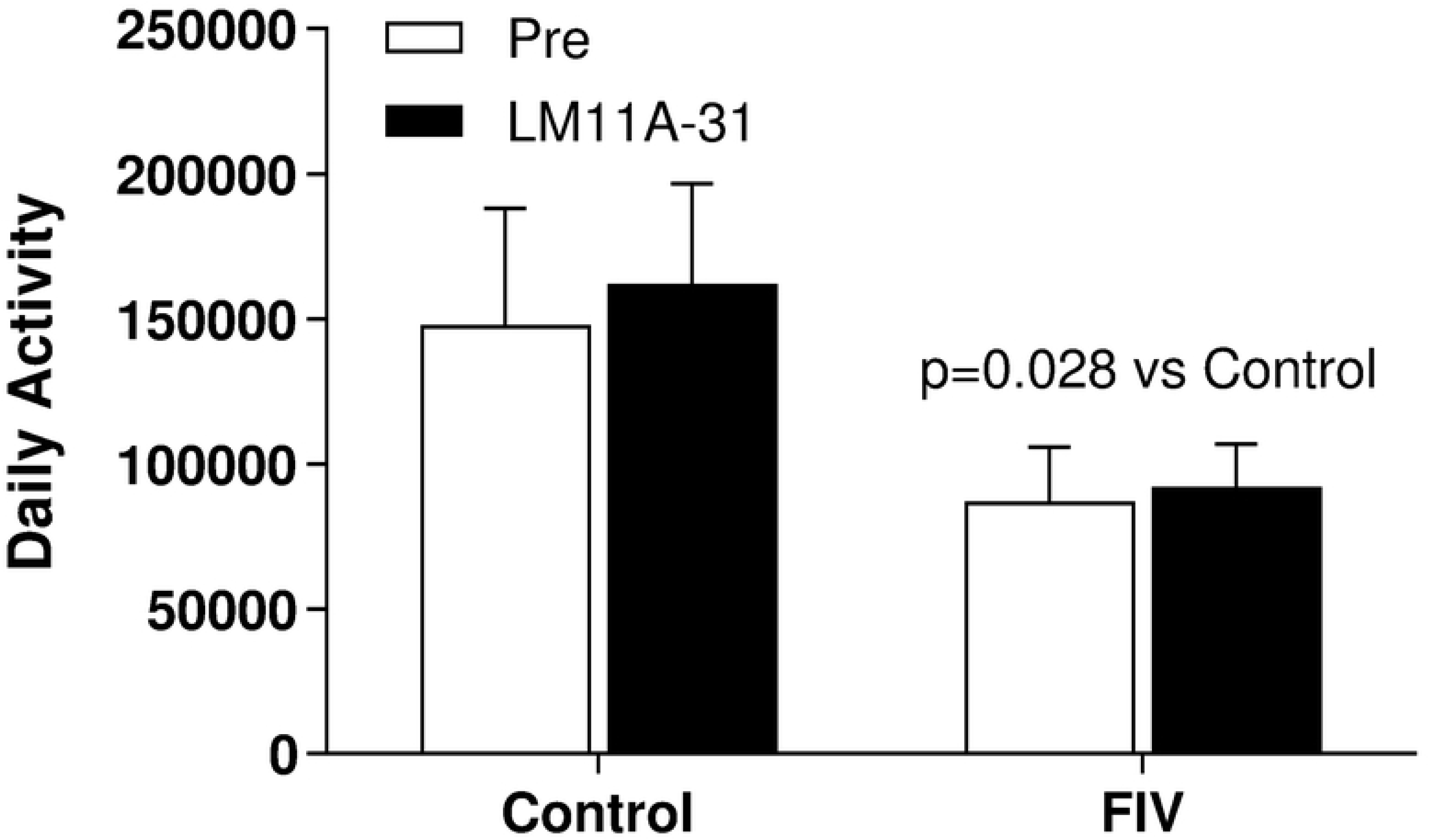
Average daily Actical monitor counts of sham inoculated control cats and FIV infected cats prior to and during treatment with LM11A-31. FIV infected cats were less active than uninfected (FIV (n=6) vs Control (n=5), 2 way ANOVA, p=0.028). No significant changes in total activity were seen in response to LM11A-31.

### T maze performance improved with LM11A-31 treatment

By 12 months post inoculation, the FIV infected cats began to show increased and more variable latencies in the reversal and repeat reversal sessions of the T maze. To evaluate the progression of cognitive decline from 12 months to 18 months, the changes in latencies were compared during discrimination, reversal and repeat reversal in the sham controls, FIV and FIV+LM11A-31 groups. The average trial latency for the first 4 sessions (which all cats performed) at each test point are illustrated for sham controls, FIV and FIV + LM11A-31 in Figure 7A. Uninfected sham control cats generally showed stable latencies after the first trial with gradual improvement over sessions. By 18 months, the average latency times for the control group were consistently low. In contrast, all FIV infected cats increased reversal and repeat reversal latency times with less stability at 18 months relative to 12 months. Taken together, these data clearly indicated the onset of a decline in performance. FIV cats receiving placebo continued to show decreased performance (increased latencies) at 20 months. In contrast, FIV-infected cats treated with LM11A-31 showed response patterns that were similar to controls, with low latency times and decreased variability. The mean change in latency illustrates the improvement in the LM11A-31 treated cats (negative differential latencies) relative to the placebo treated cats (increased latencies, Fig. 7B). We utilized an additional crossover session to determine if the placebo FIV cats would improve performance following treatment with LM11A-31. A similar pattern of improvement was observed in the crossover group when compared to their latencies while under placebo, further strengthening the evidence for a positive drug effect (Fig. 7C). A comparison of the number of errors made at 18 months relative to performance at 12 months (Fig. 7D) showed a small improvement for the LM11A-31 cats (decreased errors) whereas the performance of both the sham and FIV cats declined slightly (increased errors). The trend toward improved performance in the treated cats failed to achieve significance (p=0.08) but was consistent with the improved latencies. When the ability of the cats to learn the discrimination and reversal paradigms was evaluated by comparing trials to reach criterion, no significant differences were seen, indicating that all cats were able to learn the task equally well (Fig 7E).

**Fig 6.**
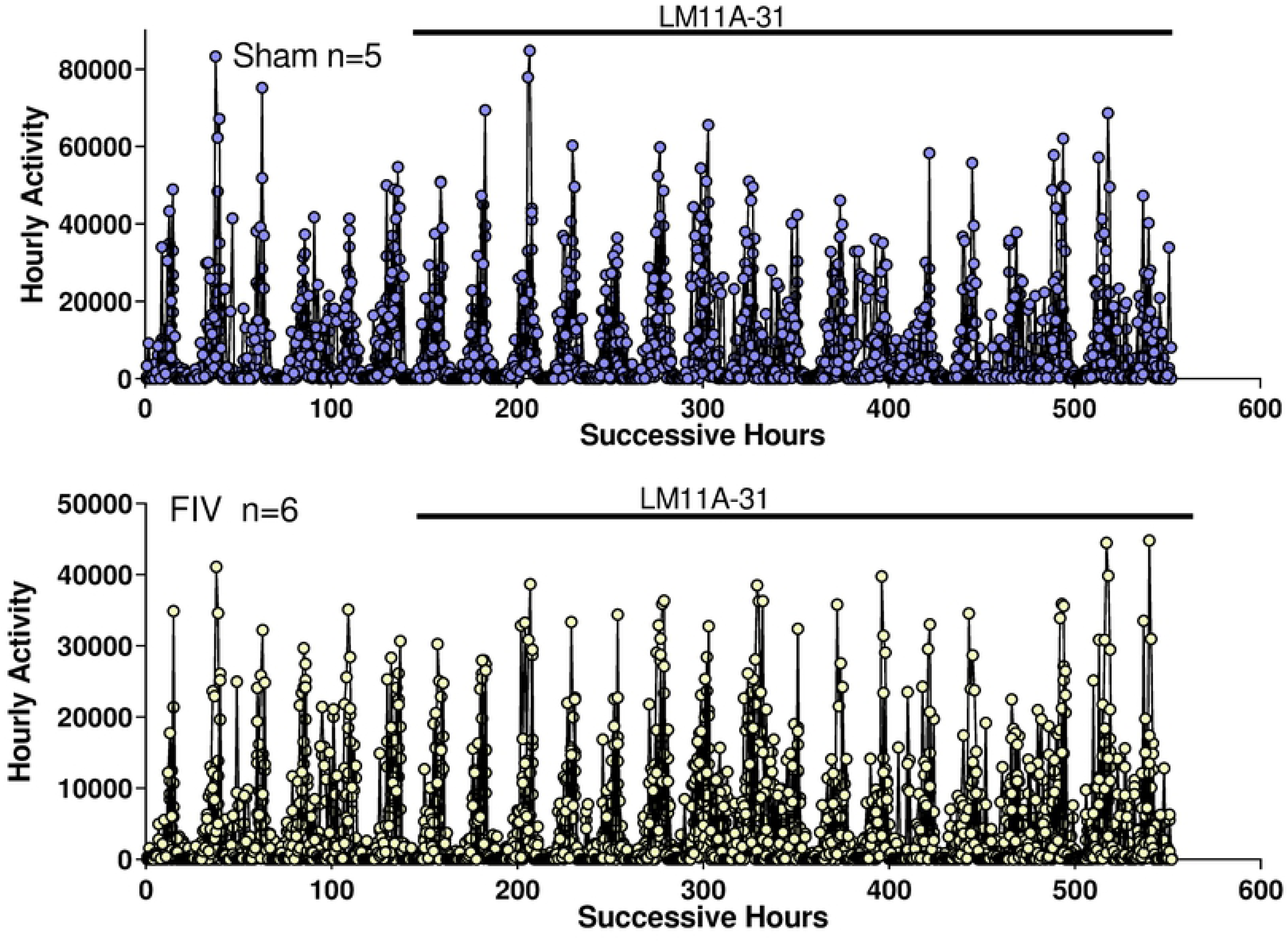
Hourly activity patterns of sham inoculated (n=5) and FIV infected cats (n=6) before and during treatment with LM11A-31. No significant changes in activity patterns were detected.

**Fig 7.**
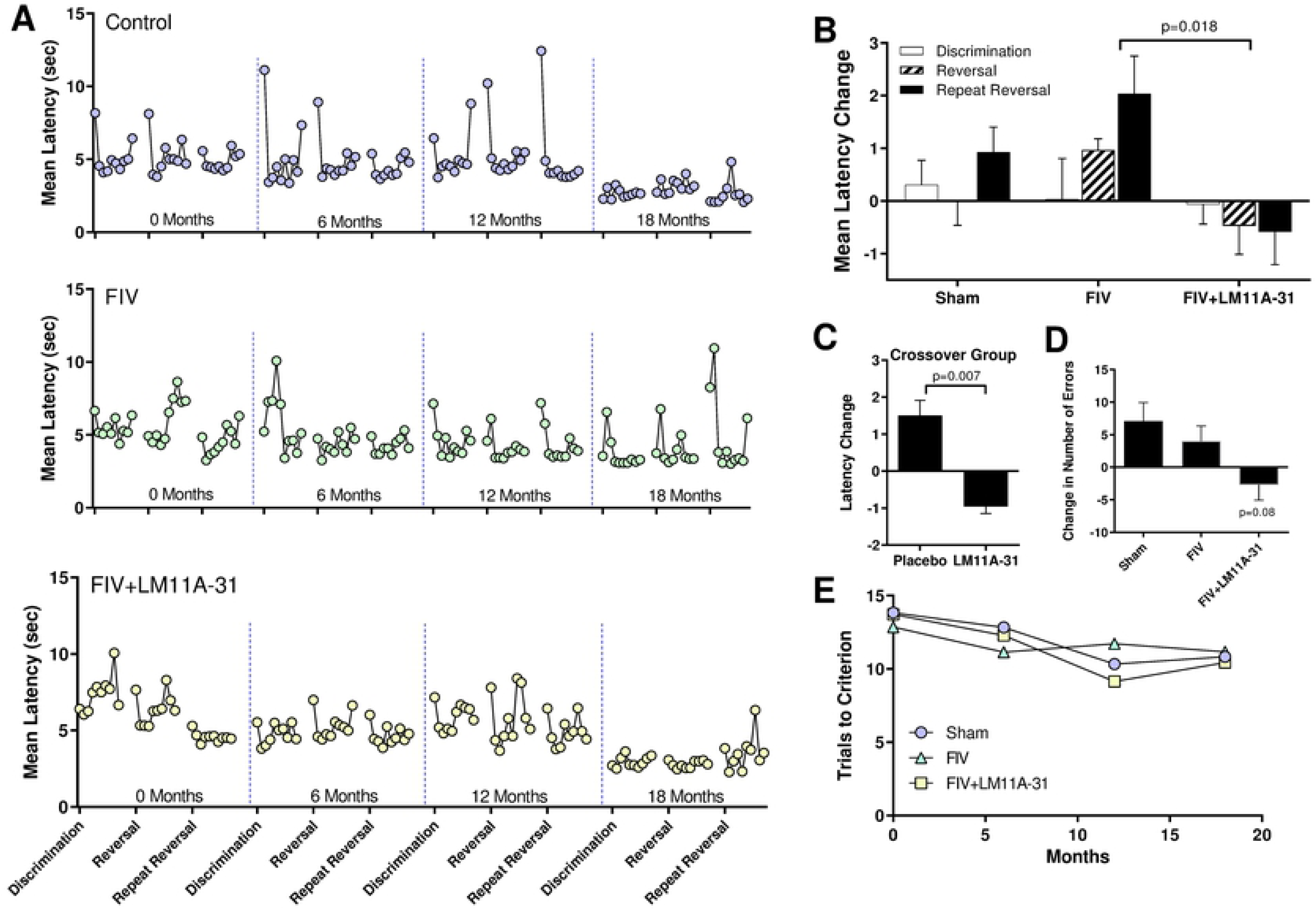
LM11A-31 treatment improved T maze performance. A. Average trial latency times over the first four sessions of discrimination, reversal and repeat reversal testing at 0 (pre-inoculation), 6, 12 and 18 months after FIV inoculation. During treatment with LM11A-31 (yellow circles, lower), latencies decreased and became much more stable, resembling sham inoculated control cats (blue circles, top). High first trial latencies for many sessions of the sham control cats were due to one cat that often showed a high latency on the first trial only. B. Analysis of the change in mean latencies after LM11A-31 treatment versus before treatment illustrating the increase in reversal latencies seen in the FIV infected cats in contrast to the decrease in reversal latencies in the cats treated with LM11A-31. C. Crossover of placebo cats to treatment with LM11A-31 also showed an improvement in latencies (matched t-test, pre-treatment vs. post-treatment, p=0.007). D. A slight reduction in the average number of maze errors per session was seen in the LM11A-31 treated cats relative to placebo controls but just failed to reach significance (p=0.08). E. No changes were seen in the number of trials required to reach criterion.

### A relationship between T maze performance and FIV titers was seen before but not after treatment

To evaluate the relationship between viral titers and T maze performance, we correlated CSF, plasma and PBMC FIV titers with mean reversal latencies. A significant positive correlation was seen between pre-treatment plasma FIV (r=0.519, p=0.027) and latencies (Fig. 8A). A trend was also observed between CSF FIV and latencies (r=0.421, p=0.082; Fig. 8B). Finally, PBMC proviral burden also exhibited a positive correlation with latency (r=0.640, p=0.0039; Fig. 8C). Since CSF FIV decreased in response to treatment, we then asked if the improvement in latencies during treatment might be related to the observed decreases. No relationship between the change in CSF FIV titers and improvements in latencies was noted (r=0.193, p=0.593, not shown). Plasma and PBMC FIV measured after treatment did not correlate with the change in latency (r=0.340, p=0.306 and 0.255, p=0.449, respectively, not shown) suggesting a potential loss of the impact of the virus on performance.

**Fig 8.**
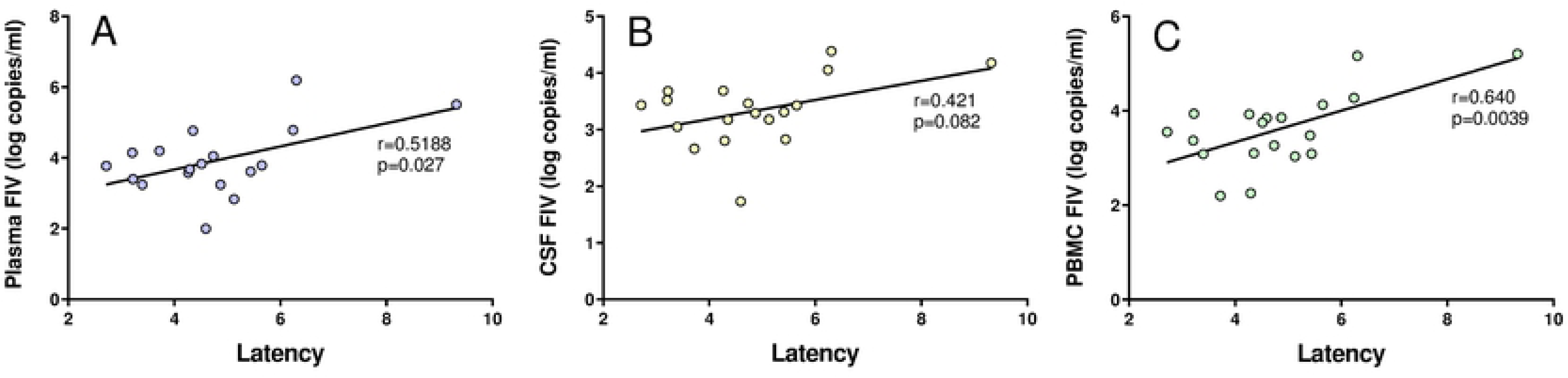
The relationship between FIV RNA copies and T maze latencies at 12 months. A. A significant positive correlation was seen between average latencies and pretreatment plasma FIV (r=0.519, p=0.027). B. A similar trend was noted between pretreatment CSF FIV and latencies but did not reach significance (r=0.421, p=0.082). C. Pretreatment PBMC FIV DNA also correlated with latencies (r=0.640, p=0.004).

### Improvements in latency in response to LM11A-31 treatment correlated with greater MAP-2 immunoreactive pyramidal cell and Iba1 positive microglia density

To determine if latency increases in the FIV infected cats were associated with neural damage, we evaluated expression of microtubule associated protein-2 (MAP-2). Loss of MAP-2 immunoreactivity is a sensitive marker of neural damage that often appears before other signs of disease (20, 26, 54–56). Since in vitro studies have shown that LM11A-31 preserves MAP-2 under pathological conditions, we hypothesized that we would see a relationship between improvement in performance and the post-treatment levels of MAP-2. Postmortem staining for MAP-2 was performed for 21 of the cat brains. Contrary to our hypothesis, regression analysis of MAP-2 immunoreactive pyramidal cell density in the hippocampus showed no correlation with average latencies (not shown). We then asked if the MAP-2+ hippocampal pyramidal cell density correlated with improvements in performance rather than absolute latencies (i.e. a decrease in average T maze latency) from 12 to 18 months. A significant negative correlation was seen when pyramidal cell density in hippocampal regions CA1 (Fig. 9A) and CA3 (Fig. 9B) was compared to the change in reversal latencies, suggesting that improvements in performance were directly related to hippocampal neuron damage. A similar comparison of Iba-1 immunoreactive microglia to latency change was also performed to examine the relationship between performance changes and inflammatory activity. As shown in Figures 9C and 9D the relationship was the opposite of the predicted outcome. In the hippocampal dendritic field (stratum radiatum and lacunosum), greater numbers of Iba-1 immunoreactive microglia were associated with improved latency scores (negative change scores, r=-0.580, p=0.019). No relationship was seen between the Iba-1 immunoreactive microglia in the pyramidal cell layer (stratum pyramidale) and the latency changes (r=0.354, p=0.215). It is noteworthy that the overall density of Iba-1 immunoreactive microglia in the hippocampus was relatively low as compared to results from more aggressive animal models of disease, indicating that the extent of inflammation was modest.

**Fig 9.**
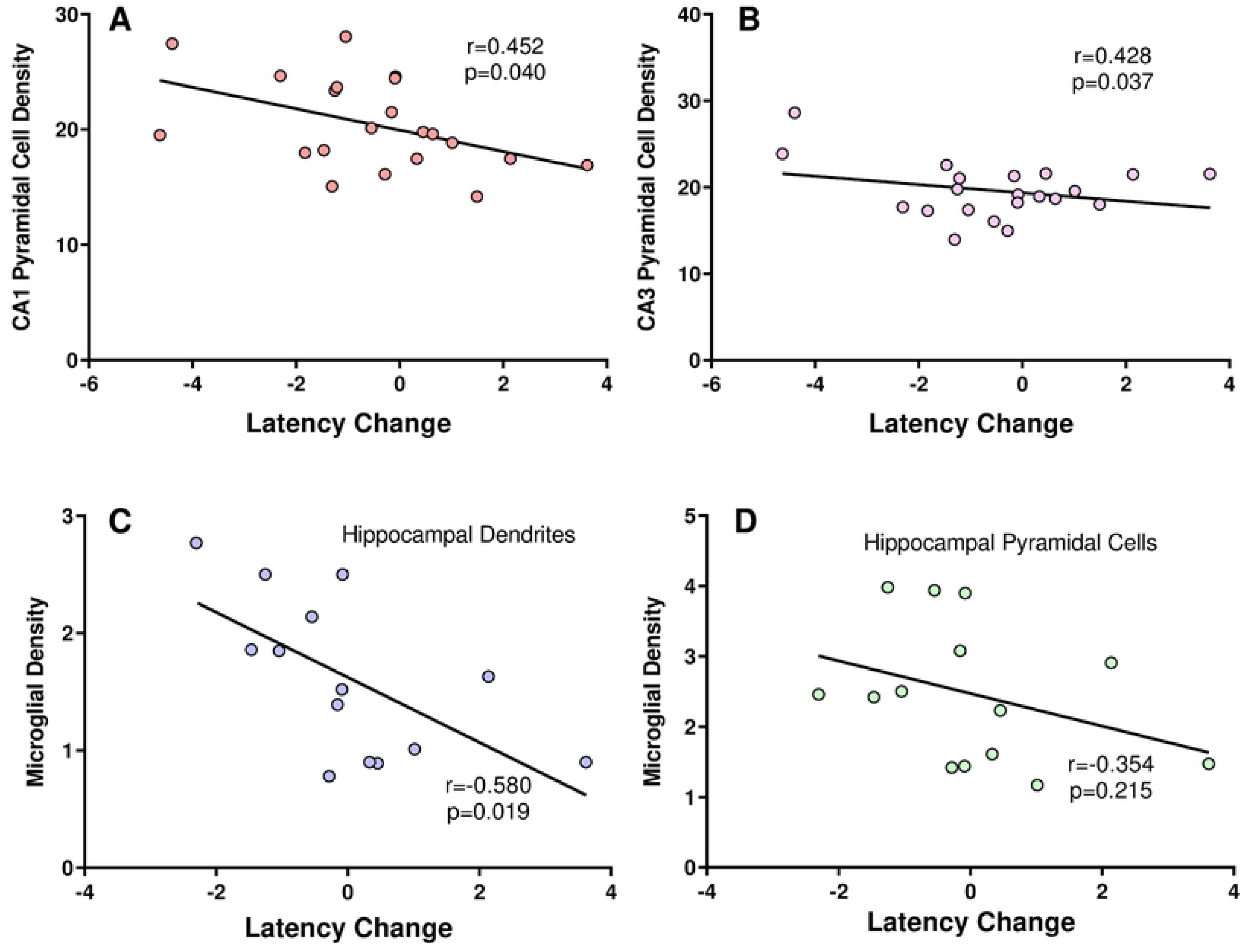
Correlations of neuronal structure (MAP-2 positive stain) and microglial activation (Iba1 stain) with decreases in T maze performance. Decreased MAP-2+ pyramidal cell density (fluorescence brightness units) in CA1 (A) and CA3 (B) correlated with increased latency (poorer performance) in the T maze. C. A negative correlation was seen in the comparison of Iba1 positive microglial density in the pyramidal cell dendritic field and changes in latency from the 12 to 18 month test period. D. No significant relationship was seen between Iba1 stained microglia in the pyramidal cell layer and change in latency.

### LM11A-31 treated cats sustain their ability to recognize a novel object

All cats showed strong novel object recognition at all tests. By 12 months 97% of the time spent exploring the two objects in the field was devoted to the novel object with a range of 11.2 – 27.7 sec. Exploration of the familiar object was relatively stable with an average range of 0.6-1.6 sec. By 18 months, we recorded a large drop (45%) in time spent exploring the novel object for the placebo FIV infected cats. In addition, 47% of the FIV placebo cats showed a slight increase in the time spent with the familiar object (i.e. less familiar object recognition) which was just short of significance (p=0.086). In contrast, FIV infected cats treated with LM11A-31 demonstrated an increase relative to the pre-treatment baseline level (Figure 10). Analysis of time spent with the novel object relative to pretreatment (12 month) levels indicated a significant improvement in the LM11A-31 treated cats relative to placebo controls. These results suggest that treatment with LM11A-31 preserved the ability of the cats to recognize the novel object.

**Fig 10.**
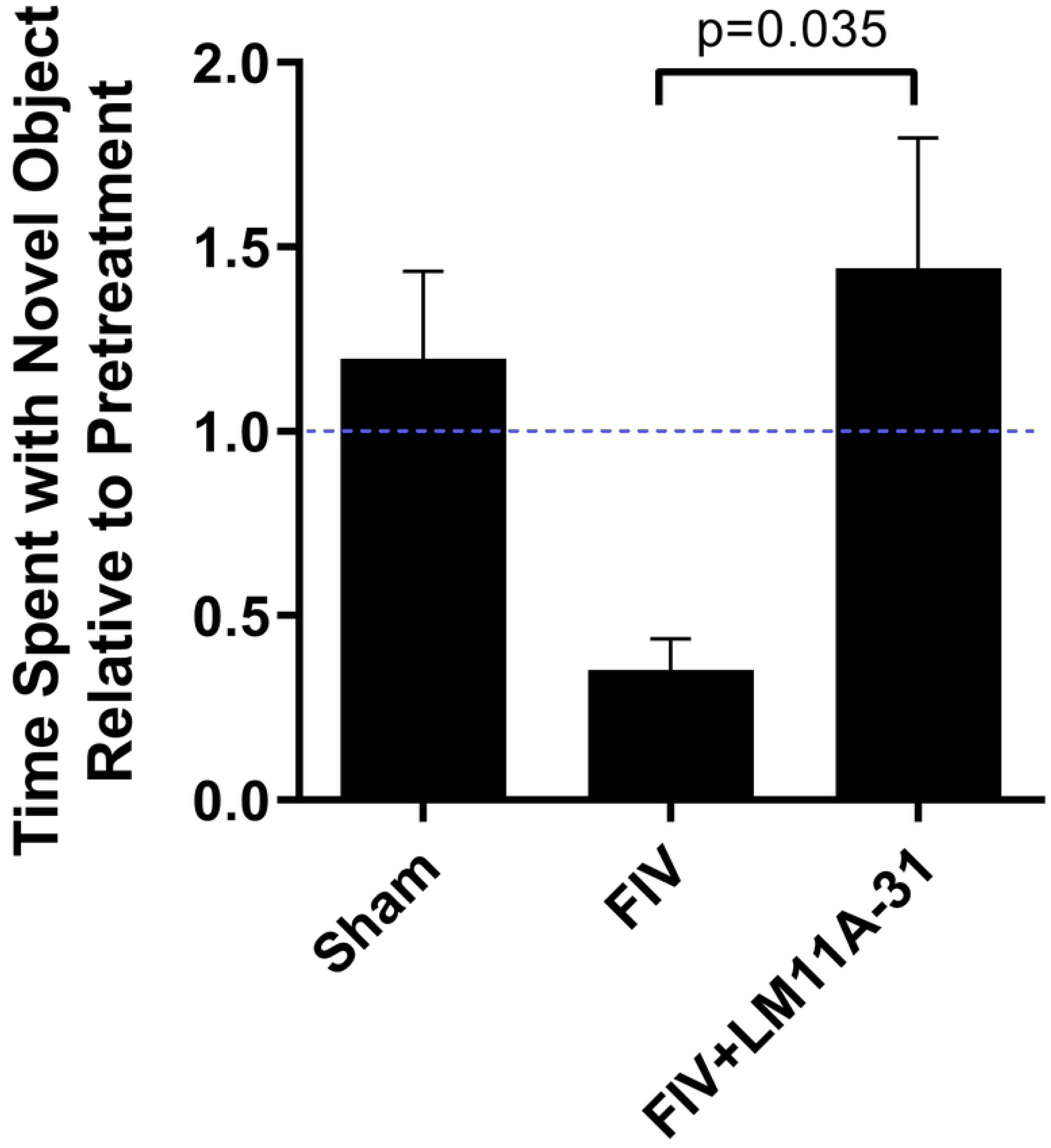
Time spent with a novel object relative to pretreatment. FIV infected cats treated with LM11A-31 (FIV + LM11A-31) retained their ability to recognize a novel object while placebo treated cats (FIV) showed a decrease in time spent with the novel object. A significant effect of LM11A-31 was seen in comparison to placebo (unpaired t-test of FIV + LM11A-31 vs. FIV p=0.035)

#### LM11A-31 treatment decreased escape duration in the open field relative to placebo

Measures of general activity (total activity, thigmotaxis) and special preference (relative time in center) in the open field were similar in all cats. FIV infected cats exhibited an increasing desire to “escape” the open field from 12 to 18 months post-infection (Figure 11) based on activity at the door (escape duration) and time spent near the door (door duration). This contrasted with the uninfected cats, which showed a decrease in escape and door duration over time, indicative of habituation to the open field environment. Treatment with LM11A-31 resulted in behavior similar to controls where the “escape” behaviors during treatment decreased relative to the pre-treatment measures at 12 months. Comparison of the behavioral changes in placebo versus LM11A-31 treated cats showed a significant improvement in response to LM11A-31 treatment.

**Fig 11.**
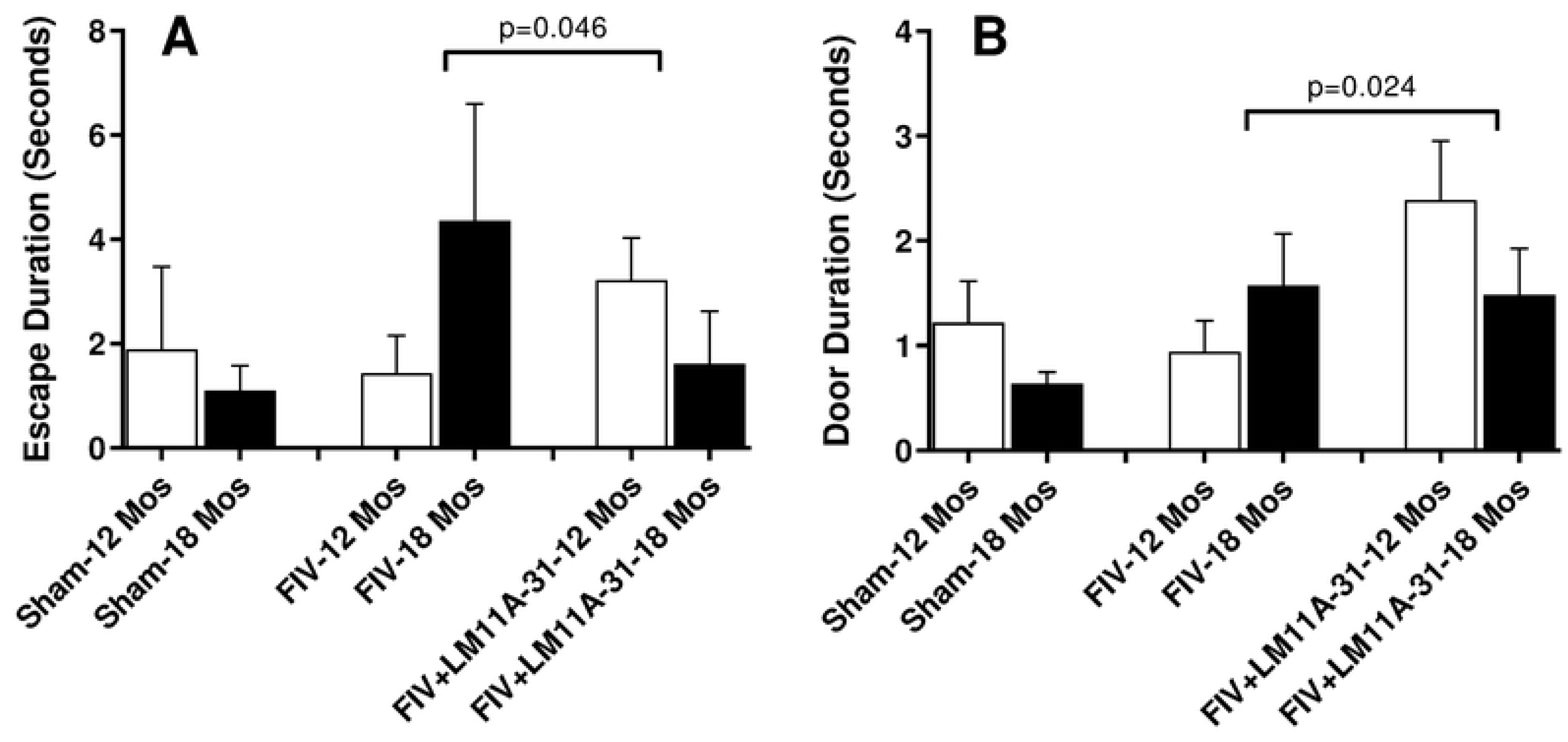
Open field escape and door duration for sham, FIV and FIV+LM11A-31 groups at 12 month (pretreatment) and 18 month tests (during treatment). Comparison of the pre-treatment versus post-treatment times for the LM11A-31 cats vs. placebo demonstrated decreased times for both escape and door duration (unpaired t-test for treatment associated change with LM11A-31 vs. placebo).

### All cats performed equally well on the GTA reaching task

All cats performed well on the motor reaching task and efficiently retrieved the reward except at the longest distances. No significant differences between LM11A-31 and placebo groups were observed in the latency and efficiency of reward retrieval. The task appeared to be easy for all cats and any minor differences were most likely due to variability in the natural reaching limits of individual cats. These observations indicated that neither infection nor drug treatment had any effect on basic motor performance for a food reward.

### No significant changes were observed between groups with administration of sensory testing

Neurotrophin receptors can play a role in the development of aberrant sensory processing including the facilitation of pain syndromes. Therefore, we asked if LM11A-31 administration might affect sensory processing in FIV-infected cats. Cats were tested for sensitivity to both thermal and tactile stimuli. Average thresholds for response to von Frey stimuli and latency for withdrawal in response to a thermal stimulus are summarized in Figure 12. Each group of cats showed similar responses to the von Frey device that were relatively stable over time. Similarly, the average response to the thermal stimulus was the same for all groups with no significant change over time. Since there was considerable individual variation in these measures, we asked if a subset of cats might show changes in response to FIV infection and/or drug treatment and found no evidence for changes that might suggest any unique individual sensitivity to the drug. Thus, neither FIV infection nor drug treatment produced any significant changes in sensory perception even when assessed for each cat individually.

**Fig 12.**
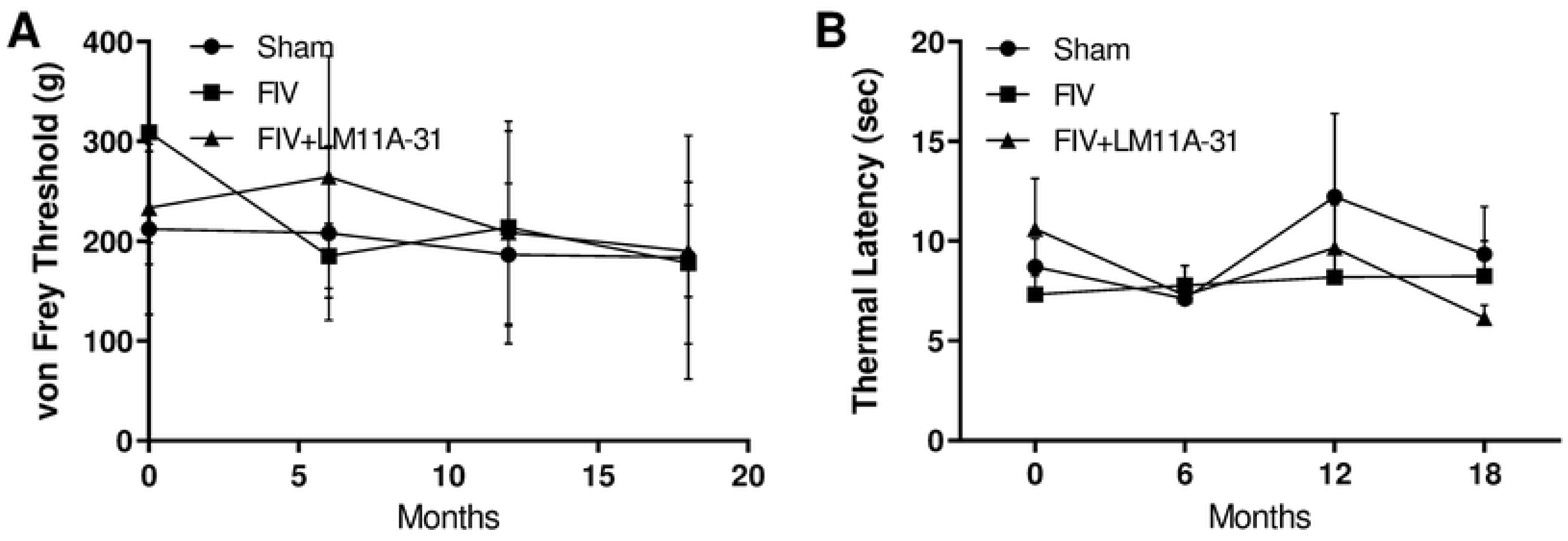
Average latency of Sham, FIV and FIV + LM11A-31 groups in response to a sensory and thermal stimulus at 0, 6, 12 and 18 month test periods. A. No significant differences in von Frey thresholds were noted over time. B. Thermal latencies were relatively stable across the experimental test period with no significant changes observed in response to time or treatment.

## Discussion

There are currently few reliable interventions proven to prevent progressive HIV-associated neurodegeneration in response to HIV in the nervous system. Studies report that neurotrophin supplementation is likely to offer significant neuroprotection, but the development of effective treatment strategies has been difficult(57). The small, non-peptide p75^NTR^ ligand, LM11A-31, mimics the beneficial effects of neurotrophin signaling and restores neural function in models of HIV associated neurodegeneration in vitro(20, 26). The compound is orally bioavailable and crosses the blood brain barrier with high efficiency making it an excellent candidate for treatment of HIV-Associated Neurocognitive Disorders (HAND). To test the safety and efficacy of the compound in an infectious animal model with gradual neurodegenerative changes, we treated FIV infected cats with LM11A-31 during chronic infection at a time when neurocognitive changes were just beginning to appear. We found that LM11A-31 had beneficial effects on the behavior of FIV infected cats, in the absence of any significant adverse effects. In addition, it also had modest but beneficial effects on virus production and T cell counts. These data are in alignment with studies from other neurodegenerative disease animal models that have demonstrated LM11A-31’s neuroprotective effects (29, 32, 34, 35, 37, 58–60).

The pharmacokinetic profile of the compound in cats, with high CSF penetration and an estimated half-life of 1.3-2.1 hours, was similar to studies in mice that showed rapid and efficient penetration of the brain with a half-life of 3-4 hours in brain tissue(32). In vitro studies have shown that the compound has nanomolar efficacy, which supports neurite outgrowth and protects from neural damage in models of disease, infection and injury(20, 26, 29, 31, 33, 34, 60, 61). The protection does not appear to be dependent on TrkA expression indicating that specific modulation of the p75^NTR^ can have a direct pro-survival impact, in part, through facilitation of Akt signaling and suppression of pathways that lead to damage(33). These effects stabilize intracellular calcium and the cytoskeleton(20, 26) which are early pathological events in a number of neurodegenerative models(21, 62–66). In vivo, these protective effects have translated to improved cognitive behaviors in mouse models of AD and accelerated recovery from spinal cord injury; while the studies here indicate improved performance in cats with CNS FIV infection. The stability of the systemic viral titers and the PBMC proviral load during drug administration alleviate concerns regarding the potential enhancement of viral synthesis via alterations of neurotrophin signaling. Although further investigation is needed, the reduction in CSF virus suggest that changes in microglial/macrophage phenotype may restrict viral replication. Since much of the CSF virus is likely to be of systemic origin(67, 68), which was not affected by treatment, the specific impact on CNS viral replication may be greater than indicated by the log difference in CSF FIV. Consistent with this possibility was the observation by Samah, et al.(69) that NGF increased monocyte chemotactic activity but decreased virus production. This may be due to a neurotrophin mediated phenotypic shift(70, 71) that is less favorable to viral replication.

A shift in the macrophage/microglial phenotype may have additional benefits by restricting the ability of the cells to mount a neurotoxic response. In vitro treatment of monocyte-derived macrophages with nerve growth factor (NGF) shifts the cells to a less neurotoxic phenotype whereas pro-neurotrophins (proNGF), which bind p75^NTR^ with high affinity, enhance secretion of neurotoxic factors(70, 71). Interference with the negative effects of the pro-neurotrophins by LM11A-3, may provide unique “anti-inflammatory” effects that offer protection. Immune regulatory effects may extend to systemic immune cells, such as contributing to the improved CD4^+^:CD8^+^ ratio in the treated cats. Although the main purpose of the T cell analyses was to show that the compound did not have deleterious effects, the small improvement in the CD4^+^:CD8^+^ ratio in treated cats suggests that LM11A-31 may have some beneficial effects upon T cell immune homeostasis.

One concern with modulating neurotrophin signaling was the potential to have adverse effects on sensory perception. The effects of NGF on pain signaling have been widely explored; therefore we were aware that NGF-like effects of LMA-11A-31 treatment might possibly enhance sensitivity to thermal and mechanical stimuli(72–75). On the other hand, studies have shown that NGF signaling through TrkA may reduce pain associated with existing nerve damage including damage in response to HIV Vpr(76). Importantly, recent studies have indicated that the beneficial effects of NGF may be dissociated from the pain sensitizing effect(77) and that the pain component may be associated with p75^NTR^ signaling(78). Actions at microglia may play a role in the beneficial effects of a mutant form of NGF(77). Thus, either sensitizing or protective effects of LM11A-31 could be possible. We used two reliable measures of sensory perception, the digital von Frey and thermal sensitivity to assess changes in the infected and treated cats. In our model we saw no significant changes in sensory perception due to FIV infection or treatment with LM11A-31. Since no evidence for a peripheral neuropathy was present in the cats, it is unclear if the compound might provide protection. It is equally important that the compound alone had no sensitizing effect on sensory perception in the absence of infection. This finding is consistent with a lack of effect on thermal sensitivity in a mouse model of AD(32). Thus, LM11A-31 does not share the adverse effects of NGF and could potentially have beneficial effects through its interactions with the p75^NTR^.

The studies were designed to initiate treatment during the development of mild cognitive impairment, in an effort to approximate the current status of HIV infected individuals. Because cats have less robust neuropathology relative to humans, the FIV was directly inoculated into the cerebral ventricles enhancing the likelihood and prevalence of behavioral changes during the 18 month study window. This strategy was successful since the intracranial inoculation resulted in deficits in most cats; whereas, based upon previous studies, many fewer cats would be expected to have CNS effects with systemic FIV infection(79). Although the intracranial inoculation speeds up pathogenesis and increases the proportion of cats that develop CNS deficits, it is important to note that the cats do not develop encephalitis, but have gradual CNS disease progression. In addition, infection within the CNS was enhanced, but not uniquely compartmentalized. A robust systemic infection was rapidly achieved, consistent with drainage of infectious virus into the cervical lymph nodes. The data were consistent with previous studies showing that the intracranial route was not only more efficient at inducing higher and more sustained CSF viral titers, but also produced consistently higher FIV titers in plasma relative to systemic (i.p.) inoculation (80). As expected, the cats in this study began to show behavioral deficits as early as 12 months post-inoculation, although the magnitude of the deficits were modest. Motor performance was quite good throughout the study based on Actical monitoring, open field activity and T maze and GTA performance. Stable body weights and body condition scores indicated that the cats were in good general health throughout the study.

T maze deficits were restricted to increased latencies and more variable performance particularly during reversal testing, consistent with impaired information processing. The increase in latencies in the FIV infected cats was less than anticipated, partly due to testing at an early stage of CNS disease and the ability of the cats to run the maze very rapidly, even with obstacles designed to slow progress (high hoops in the pathway). The measurement of latencies in the T maze was similar to time based tasks used for neurocognitive assessments in HIV infected humans(81). The modest deficits at 18 months likely reflect early stages of cognitive decline and the gradual progression of disease similar to people on antiretroviral therapy living with HIV. The significant improvements after a relatively short course of treatment are consistent with the ability of LM11A-31 to stabilize cognitive decline, as seen in other animal models of neurodegenerative disease. In some cases, the behavior normalized to the level of uninfected cats, suggesting potential reversal of deficits. Although a direct assessment of effects on CNS pathology could not be performed with the current paradigm, the correlation of latency changes with a marker of neural degeneration, MAP-2 loss, suggests that the improved latency scores may be related to stabilization of the neuronal cytoskeleton. Other studies of cognitive function in mice have shown that LM11A-31 improves performance in mouse models of AD(32, 34) and Huntington disease(36). Improvements included decreased latencies in a Y maze(32) improved novel object recognition(82) decreased latency to find the platform in the water maze(34) and improved place learning(58). In addition, evidence for enhanced restoration of neural function was seen after LM11A-31 treatment in a model of traumatic brain injury(35), spinal cord injury(37, 83) and in normal aging(59). Collectively, these studies provide strong support for broad neurocognitive stabilizing effects of LM11A-31.

The negative correlation seen between the hippocampal density of Iba1 positive microglia and latency change is more difficult to explain. In the absence of pretreatment measures of microglial density it is difficult to know the exact effect of LM11A-31 on microglial activation. Other studies in mouse models of Alzheimer disease and Huntington disease, and our data utilizing gp120 transgenic mice (Meeker, et al., unpublished data), have documented that treatment with LM11A-31 decreases microglial activation(84, 85). For the cats in the study reported here, the level of microglial activation was modest and likely reflects the early stages of inflammation or perhaps a reduction by LM11A-31. It is also important not to rule out the possibility that microglia have protective functions, particularly during the early stages of disease. The ability of LM11A-31 to shift phenotype could also offer some protective benefits not specifically linked to microglial numbers.

Novel object recognition by the FIV cats also declined at the 18 month test indicative of cognitive deficits. Beneficial effects of LM11A-31 again suggest improved cognitive function similar to the results for novel object recognition in a mouse model of AD(82). The effect was largely due to the stabilization of novel object recognition versus a decline in this measure in the placebo cats. Thus, these results are consistent with an effect that prevents disease progression.

The open field was designed to assess general motor activity, exploration and anxiety through the measurement of total movement, activity around the edges of the maze (thigmotaxis), time in the center of the maze and activity concentrated in the vicinity of the door (desire to escape). It has been widely used in rodents to test the effects of anxiolytic compounds(86) and has been successful for efficacy evaluation of some drugs (e.g. benzodiazepines) but less successful for others (e.g. SSRIs). Uninfected cats explored the maze freely and showed the expected habituation to the maze over time, including less preoccupation with exiting the maze (time spent next to the door and escape-like behaviors). The FIV infected cats in our studies showed an increased duration near the door by 18 months suggesting an increased desire to leave the maze. Cats treated with LM11A-31 at 18 months contrasted with the untreated FIV infected cats, showing a decrease in time spent near the door during drug treatment that was similar to the behavior of uninfected cats. This suggested a decrease in anxiety and was consistent with the effects of the compound in other open field studies and measures of anxiety-like behavior. Previous studies in a mouse model of sepsis-induced cognitive impairment showed that LM11A-31 normalized exploratory behavior in the open field(87). Similar results were seen in a mouse model of Huntington disease that showed a reduction in anxiety in the light:dark test following treatment with LM11A-31(36). Collectively, these findings provide clear evidence that LM11A-31 exhibits beneficial effects by ameliorating cognitive decline in an animal model for FIV neurodegeneration, while no adverse effects were associated with therapy.

### Summary

The goal of this study was to establish safety and neuroprotective efficacy of LM11A-31 in an animal model of active lentiviral infection. The intracranial inoculation, designed to optimize infection of the CNS, resulted in findings consistent with a primary infection of the CNS. Progression of cognitive changes was slow, as anticipated, and similar in many respects to the gradual progression of neurocognitive disease in individuals infected with HIV. Treatment with LM11A-31 at a point where the cognitive deficits were just becoming apparent normalized deficits in T maze performance, novel object recognition and open field behavior. Potential adverse effects such as changes in sensory perception and increased systemic viral burden were not observed with LM11A-31 therapy. A decrease in CSF FIV titers and a slight improvement in the CD4:CD8 ratio suggested that LM11A-31 may have beneficial effects beyond the anticipated neuroprotective effects. These findings are similar to beneficial effects seen in other animal models of neurodegeneration and CNS injury and support the use of LM11A-31 as an adjunctive neuroprotective agent for the treatment of HIV infected individuals.

## Materials and Methods

### Subjects

A total of 30 specific-pathogen-free, purpose-bred, neutered male domestic short hair cats (*Felis catus*) between 1-3 years of age were used. The cats were maintained in individual cages (188 cm high, 147 cm deep, 91 cm wide) in a laboratory animal facility on a 12/12 hour light–dark cycle, fed a measured balanced feline dry ration and maintained throughout at body weights consistent with initial body weights and a lean (3/9-4/9) body condition score, as referenced on a standard score chart (Purina Body Condition Score Index, http://www.purina.com/cat/weight-control/bodycondition.aspx).

### Infection with FIV

With systemic infection, neuropathological changes are slow and typically develop in approximately 20% of infected cats. To accelerate testing with the feline model we have developed an intracerebroventricular inoculation, described previously (45). This procedure increases the CSF/plasma FIV ratios and the proportion of cats with CNS disease. These cats develop mild cognitive deficits after about 18 months, similar to the deficits currently seen in HIV infected individuals on combination antiretroviral therapy.

Specific pathogen free cats were purchased from a supplier (Liberty Research, Waverly, NY) at 6 months of age and maintained in the colony at the NCSU College of Veterinary Medicine. All procedures were approved by the NCSU IACUC #15-157-B and were performed in accordance with the NIH Guide for the Care and Use of Laboratory Animals and NIH Guidelines for the Ethical Conduct of Research. Cats were induced for anesthesia in an isoflurane inhalation induction chamber, intubated with an endotracheal tube, and maintained on isoflurane inhalation anesthesia for the duration of the procedure. Following the induction of anesthesia with isoflurane, cats were inoculated intracranially into the right lateral ventricle with a combination of FIV_NCSU1_ and FIV_V1CSF_. All inoculations were performed under sterile conditions in a dedicated veterinary surgical suite at the College of Veterinary Medicine. An approximate 2 mm diameter opening in the skull was created with a high-speed dental burr. A custom 23- gauge stainless steel cannula connected to silastic tubing filled with sterile artificial CSF and backloaded with 250 µl of FIV (10^5^ TCID_50_ FIV_NCSU1_/FIV_V1CSF_ per 200 µl) was inserted into the brain 3.5 mm lateral to midline at AP 14.5 mm to a depth of 10 mm below the surface of dura. The silastic tubing was elevated to a height of approximately 50 cm, unclamped and the cannula was slowly raised (if necessary) until fluid flowed freely into the ventricle. After delivery of 200 µl, the tubing was clamped. The cannula was left in place for 2 minutes to allow diffusion of the virus. This procedure allowed delivery of virus under minimal pressure to prevent disruption of the blood brain barrier or damage to brain tissue. The cannula was then slowly raised out of the brain and any signs of blood or reflux of CSF noted. Bleeding from brain tissue was not observed in any cat, and reflux of CSF was negligible. The opening in the cranium was sealed with bone wax and the incision was closed with buried absorbable sutures to minimize postsurgical complications from the wound (e.g. scratching). The cats were given a dose of buprenorphine (0.02 mg/kg, sublingually) and closely observed during recovery. The cats were alert and ambulatory within an hour post surgery and minor complications such as mild incisional discharge, were rare. Postmortem analysis of the brain at the end of the study showed no signs of damage in the brain and skull for any of the cats, indicating full recovery from the surgery and no CNS trauma. Uninfected control cats underwent the same surgical procedure without virus inoculation, to control for any effects of the surgical procedure and to insure that investigators were blinded regarding which cats were infected. To verify that the cats appeared healthy and normal, the testers and veterinary technicians, who were blinded to treatment conditions, were surveyed at the end of the study to indicate which cats they thought were FIV infected. The average responses were at the chance level (51% correct for infected vs not infected) and randomly mixed indicating that personnel could not discriminate between the infected and uninfected cats.

### LM11A-31 treatment

Cats were treated with a pharmaceutical grade salt of LM11A-31 (LM11A-31-BHS) identical to the compound currently under investigation in human trials. The dosage was selected based on data from previous mouse studies, which demonstrated efficacy at a single daily dose of 50 mg/kg. The dose was adjusted for cats based on body surface area and was validated in initial pharmacokinetic studies. Although previous studies showed efficacy with once daily oral administration, twice daily dosing was chosen to maintain more consistent drug levels, given the relatively short half-life, and to match the strategy for human clinical trials. Based upon the mechanism of action, it is likely that the modulation of neurotrophin signaling has effects that greatly outlast the circulating drug; however, the long term stability of these effects is currently unknown. Oral dosing of the cats was accomplished by adding a concentrated solution of the drug or placebo (78-128 µl) to a small “meatball” of canned feline cat food that the cats would readily ingest. Cats were treated in the morning (7:00-8:00 AM) prior to testing and at the end of the day after all testing was completed (5:00-6:00 PM).

### FIV PCR

One week following surgery, while under isoflurane anesthesia, a sample of blood was drawn from the jugular vein and a sample (∼1 ml) of CSF was collected from the cisterna magna in each cat(46). The blood and the CSF were centrifuged to remove cells and a sample of each used for FIV PCR. All inoculated cats were positive for FIV and the ratio of CSF to plasma was high and consistent with a primary infection of the CNS. The inoculation also proved to be highly effective for induction of systemic infection, most likely due to rapid drainage of the virus into the cervical lymph nodes.

#### FIV provirus amplification

Primers for routine PCR amplification of FIV provirus were selected from a highly conserved gag region within the FIV genome. Primers FIV-7 (5’-TGACGGTGTCTACTGCTGCT) and FIV-8 (5’-CACACTGGTCCTGATCCTTTT) amplify an 838 base pair segment, while primers FIV-1 (5’-CCACAATATGTAGCACTTGACC) and FIV-2 (5’-GGGTACTTTCTGGCTTAAGGTG) amplify a 582 base pair segment included in the region amplified by primers FIV-7 and FIV-8. By comparison with a known amount of plasmid DNA containing the FIV gag gene (a kind gift from R. Olmsted), we can detect between 50 and 100 copies of FIV provirus using primers FIV-7 and FIV-8 and as few as 5 proviral copies using FIV-1 and FIV-2 as nested primers. All oligonucleotide primers were synthesized in-house using an Applied Biosystems DNA Synthesizer, Model 391.

For PCR amplification of provirus, PBMCs were isolated, washed 2 times with ice cold PBS, and re-suspended at 1 x 10^5^ cells/100 uL in PCR buffer with 600ug/mL proteinase K. The cells were digested at 56° C for 2 hours, boiled for 10 minutes, and stored at - 135° C. PCR amplification was performed using 25 uL of sample and primers FIV-7 and FIV-8 in previously determined optimal conditions (0.1 ug/reaction of each primer, 1.5 mM MgCl_2_, and 1.0 units of Taq polymerase). The amplification cycle of 95° C for 35 seconds, 59° C for 35 seconds, and 72° C for 35 seconds was repeated for 30 cycles, followed by a 7 minute, 72° C extension cycle. Ten microliters of amplified product was run on a 1.2% agarose gel. If a discernable amplification product was not present, 40uL of the PCR product was re-amplified for 20 cycles after the addition of 0.2 μg of the nested primers FIV-1 and FIV-2 and 1.0 U of Taq polymerase. Ten microliters of amplified product was again run on a 1.2% agarose gel. Serial dilutions of purified plasmid DNA containing the gag gene were amplified as a positive control for amplification efficiency as previously described (45, 47).

#### Real Time PCR quantification of FIV RNA and FIV DNA

Viral RNA was extracted from CSF and plasma using QIAamp® Viral RNA Mini Kit (Qiagen, CA). The viral RNA loads were determined by real time RT-PCR with the Taqman one-step RT-PCR master kit (Applied Biosystems, CA). The following specific primers and probe of NCSU_1_ FIV gag region were used: FIVNC-491F (5’-GATTAGGAGGTGAGGAA GTTCA GCT –3’), FIVNC-617R (5’-CTTTCATCCAATATTTCTTTATCTGCA –3’) and the labeled probe, FIVNC-555p (5’-FAM-CATGGCCACATTAATAATGG CGCA -TAMRA-3’ (Applied Biosystems, CA). These reactions were performed with a Bio-Rad iCyclerTM iQ and analyzed using the manufacturer’s software. For each run, a standard curve was generated from standard dilutions with known copy numbers and the RNA in the samples was quantified based on the standard curve(48, 49).

### Lymphocyte Subset Analysis

Whole blood from each cat was collected into EDTA anti-coagulant tubes at various times post infection for a complete blood count with differential and lymphocyte phenotype analysis. Percentages of CD4+ and CD8+ T lymphocytes were determined by two- and three-color flow cytometry. The fluorochrome-conjugated monoclonal antibodies used in staining were PE-conjugated anti-CD3(mAb 572, NCSU Hybridoma), FITC-conjugated anti-CD4 (mAb 30A, NCSU Hybridoma), PE-conjugated anti-CD8 (mAb 3.357, NCSU Hybridoma), and PE-conjugated anti-CD21 mAb (Biorad, CA2.1D6)). Isotype-matched irrelevant antibodies were used as controls. The percent of positively stained lymphocytes was determined using a Becton Dickinson LSR II flow cytometry unit. Approximately 20,000 gated events were acquired and analyzed using FlowJo software. Complete blood cell counts were determined by the veterinary clinical diagnostic laboratory at NCSU-CVM and the absolute lymphocyte number determined by differential counts. The number of cells within each lymphocyte subpopulation was then calculated based on the subset percentages obtained by flow cytometry and the absolute lymphocyte count reported on the CBC.

### Clinical Laboratory Analyses

A complete blood count with leukocyte differential, serum biochemistry profile and urinalysis were performed for each cat, using standard evaluation protocols, in the clinical diagnostic laboratory at North Carolina State University College of Veterinary Medicine. Values were compared to previously established values for healthy cats as well as to each cat’s pre-infection and pre-treatment baseline.

### Behavioral and physiological assessments

#### T maze design

A feline-adapted T-maze designed by CanCog Technologies constructed of plywood sealed with polyurethane to conform to laboratory standards, was used to provide a simple test of cognitive and motor ability(50). Cats were adapted to the maze as well as transport and handling by the testers prior to the start of testing(51). The T-maze was housed in a behavior testing suite within the same building where the cats were housed. None of the testers participated in invasive procedures such as anesthesia and blood collection, to facilitate positive interactions with the cats. Cats were placed in a start box with a manual sliding door opening to a runway containing a series of hoops, which the cats had to jump through. At the end of the runway, the cats had to decide to turn into the left or right reward arm, which then turned again leading back to the start area. Doors with magnetic latches positioned in each reward arm, were remotely closed after the arm choice was made to prevent path reversal. The reward area was hidden from view until the cat was in the reward arm and contained a disposable well with a food reward. At the end of each trial, subjects were able to pass directly back into the start box from either reward area, when a connecting door was raised. Fitted acrylic sheets covered the top of each section of the maze to prevent escape but allowed the animals’ behavior to be continuously observed. The tester stood behind the start box, positioned at the middle point, and could visualize the cat but did not provide cues or interact with the subject. A specific computer program (CatCog) was developed by CanCog Technologies to record the number of correct choices and latency to response (in milliseconds). The computer was positioned outside the cat’s range of view from within the box. The timer and reward arm doors were manually controlled by the experimenter. The order of the cats testing was randomized daily and all testers were blinded to both infection status and treatment condition. Inter-tester reliability by the three trained testers was evaluated during the study using video recordings of tester performance. An assessment at the end of testing verified that the testers could not distinguish which cats were infected (a chance level of 51% correct was measured in their assessment of infected vs. non-infected). To provide motivation, cats were trained and tested prior to daily feeding from 0800-1100 hours using highly palatable food rewards (Canned food (Hills A/D), Pounce® cat treats, Del Monte Foods; Whiskas® cat treats, Mars, Inc; various flavors). After testing, the cats were then fed a measured ration of dry chow based on body weight. The test protocol developed by CanCog Technologies (unpublished data) consisted of 6 stages: Adaptation, Reward approach, Preference testing, Discrimination training, Reversal 1, and Reversal 2 (described below). Cats were tested 6 days per week by technicians who were both familiar to and with the cats during the behavioral conditioning period. The maze was cleaned between sessions and left to air-dry.

#### Adaptation and reward approach

During adaptation, cats were allowed to explore the runway and both arms of the maze with small food rewards placed throughout the maze. After the initial sessions, the maze doors, including those to the start box, were opened and closed manually by the experimenter to acclimate the cats to the sound and associated air movement. When the subject moved reliably throughout the maze, from the start box to both reward arms over a ten-trial session, treats were restricted to the reward wells of both arms of the T-maze and high hoops were positioned in the maze for all subsequent trials. Two high hoops with a 21 cm diameter round opening and 40 cm height to the bottom of this opening were placed in the runway and one high hoop was placed in each arm in order to increase the motor difficulty of the task and encourage cognitive-motor delay. The reward-approach phase was completed when the cat traversed the maze successfully from start box to either reward arm, 10 times in one 10 min session, with the high hoops in place, and food rewards located only in the reward wells.

#### Preference testing

Following successful completion of the reward-approach stage, each cat had one day of preference testing to determine its preferred side. This was established empirically as the side that a cat went to ≥6 times during one session of 10 trials, when both arms contained rewards. Each cat demonstrated a preferred side, which was used as the first rewarded side in discrimination training. Using each cats preferred side improved performance consistency between cats and allowed us to establish a strong response pattern prior to the introduction of Reversal 1 and Reversal 2.

#### Discrimination training and reversal

The test cat was positioned in the start box and the tester started the software timer when the door out of the start box was opened. The timer was stopped after all four feet of the cat crossed a pre-determined point in either reward arm. The recorded values represented the latency to complete the trial. When stopped, the software began a 30-second inter-trial interval, which allowed the cat time to ingest the reward and return to the start box and for the tester to reset the rewards in the reward arms. To control for auditory cues, the tester lifted the doors on both the left and right reward arms and placed a reward into the empty reward well then closed both doors. Each cat completed a total of 10 trials (1 session) per day with the reward on only one side of the maze. Cats had 60 seconds to complete the maze or were recorded as a non-response for that trial. For analysis, a ceiling of 20 seconds (s) was placed on the latency measure to minimize skewing of the data. Two cats that performed inconsistently with numerous high latencies and distractions in the maze at all test periods (> 3 standard deviations from the group mean) were not analyzed.

To assure consistency in performance while leaving some flexibility for daily variation, cats were tested for a minimum of four days and a criterion of 27/30 correct responses on three consecutive days was used in order to advance to the next phase. This was because pilot studies indicated that an individual cat’s performance may vary from session to session. For example, 10/10 correct on one day may be followed by 8/10 correct on a subsequent day. After the discrimination training was completed, Reversal 1 was initiated by placing the reward in the opposite T-maze arm. Reversal 1 was followed by Reversal 2 with the reward returned to the original, preferred arm of the T-maze. Preferred side (left or right), number of correct responses, number of trials to criteria, and latency to reward arm (in milliseconds) were collected for each cat. The average time to completion of the T maze protocol was 16.3 days

#### Crossover session

To provide additional support for efficacy, cats in the placebo cohort which were showing signs of cognitive decline, were entered into a crossover cohort, where they were given drug for a period of 10 weeks. The performance of the cats was then compared to their pretreatment values.

### Open Field

The open field arena is an enclosure (288(L) x 212(W) x 246(H) cm) lined with corrugated PVC roof panels (Lowes 2.17’ x 12’) attached to wire kennel caging panels with zip ties. A single door was used for placement of the cat which is closed with two external nylon cords to prevent escape. The arena was housed in a room inside the same building where the cats were housed. The behavior of the cats was monitored through a black and white Panasonic WVBP330 digital camera connected to Panasonic Digital Disc Recorder WJ-HD309A with an Extension Unit WJ-HDE300. Connecting to that was a Panasonic DMR-EH75V DVD recorder with audio sourced from a Canon HDV 1080i camcorder. The video was displayed on a small RCA television, which allowed the tester to watch the cat remotely without disturbance.

Each cat was filmed in the open field for 10 minutes a day for 5 consecutive days. Testing was repeated every 6 months. The setup minimized any sounds from the tester or external sources. Vocalizations, rears, escapes and jumps were recorded during testing. Once the testing was completed, the DVD was transferred to a PC and evaluated using Ethovision (Noldus) software. Objective measurements included: distance moved and time spent in various areas including walls (thigmotaxis), door, and center. Vocalization, escape, rearing on the hindlimbs, and jumps were also recorded.

### Novel Object Recognition

Each cat was placed in the open field with 2 objects designated A and B. The behavior of the cat was automatically recorded using a video camera coupled to a computer running Noldus Ethovision software. The amount of time exploring/sniffing each object was measured in four 3- min sessions. Cats were exposed to two objects, A and B supported on pegs at opposite corners of a board set on the floor of their cage for session 1. For session 2, the board was rotated 45 degrees with A and B on same pegs to correct for any position bias. In session 3, object B was replaced with a new novel object C.

In session 4, an additional object C scented with a synthetic feline facial pheromone (Feliway, Ceva Animal Health) was used to assess exploration of a novel smell. During testing, the tester waited outside the room to prevent the cat from seeking human interaction. The time sniffing each object over a period of 3 min was recorded with a video camera set up on a tripod. An inter-trial interval of 3 min within the cat carrier was imposed between each session. The video was scored later by a single technician who recorded the number of sniffs per object and time spent sniffing.

### General Test Apparatus (GTA)

Initially cats were screened for their ability to perform discrimination learning tasks in a modified general test apparatus (GTA). Tests were conducted using previously established protocols and a technical staff experienced in conducting these tests. The tests included simple motor tasks, object discrimination learning, object reversal and delayed non-matching to position. Although the cats were able to perform the discrimination learning tasks, the reliability of the behavioral performance was too low to be of value to discriminate changes with drug intervention. A simple reaching task to pull in a reward tray initially showed the greatest potential. The cat was required to reach with its paw and pull a sliding coaster towards them to obtain the food reward. Testing was over a period of 3 days. On day 1 the baited coaster was initially set at position “1” and the distance from the cat was subsequently increased each trial by one position (2.5 cm each) until at position “12”. The computer was used to time latency to reward retrieval. A tester observed the cat to record the paw used to retrieve the reward, and paw changes (using one paw and then the other to retrieve the reward). This test had 12 trials with 30-second inter-trial intervals. On day 2, the distance from the cat was varied randomly between positions 5, 7, 9 and 11 for 12 trials followed by a distance of 12 for 3 trials, which was beyond the reach of most cats. Latency to retrieval, maximum distance achieved, and efficiency of retrieval at each distance was compared across groups.

### Sensory testing

Quantitative sensory testing was performed in a dedicated space adjacent to the housing area (5×5 meter isolated room with one door and no windows) at the NCSU College of Veterinary Medicine. Quantitative sensory thresholds were collected at baseline (prior to any procedures) and then every 6 months for the study duration using the electric von Frey and Thermal probe.

#### Electronic von Frey (EVF)

The electronic von Frey device (IITC model Almemo 2450; IITC Life Sciences Inc., Woodland Hills, CA, USA) consisted of a 1000 g internal load cell connected to modified 0.5 mm diameter pipette tip, as described previously(52, 53). The amount of force applied to a test site was measured and displayed with a resolution of 0.1g and maximum of 1000g (although the max force to be applied was set a priori at 500g).

#### Thermal probe

The thermal device (NTE-2A, Physitemp Instruments, Clifton, NJ, USA) was comprised of a hand-held probe equipped with flat 13 mm diameter tip on a 5-foot lead connected to a digital temperature control unit and recirculating pump with combined water reservoir. The temperature range of the probe tip could be altered to have a temperature between 0 to 50 °C, maintained to within 0.1 °C. This device was used to deliver a hot thermal stimulus (46.5°C).

Testing was performed at the dorsal metatarsal region on both hindlimbs. The von Frey device was applied perpendicular to the dorsal surface of the metatarsus, between metatarsal bones III and IV as previously described in dogs(52). The Physitemp probe was applied to the dorsal mid-metatarsal area following removal of the hair over a 2 x 2 cm area.

The end point for both thermal and mechanical stimuli was defined as a behavioral response indicative of a conscious perception of the stimulus: movement of the limb away from the tip/probe with conscious perception; turning head to look directly at the site; vocalization; or other consistent, clearly recognizable body movement indicating perception of the stimulus. Simple reflex movements (such as twitching) were not considered an endpoint.

#### Protocol for sensory testing

The ambient testing room temperature was maintained between 20.0-23.0°C. All cats were acclimated to the testing room for 30 minutes each day for 1 week prior to testing each month. On testing days, the cats were allowed to explore the room for 10 minutes prior to testing. Testing was always performed between 12pm and 2pm by the same individual, and gentle restraint of the cats was provided by an assistant (same assistant at all times). Gentle restraint of the cats consisted of the assistant standing beside the exam table, with one arm under the cat’s abdomen while the assistant’s other hand was used to hold the cat in position. The von Frey data were collected on one day, and the thermal latency data collected 1-3 days later. Both limbs were tested, and the order was randomized at the start of the study (www.randomizer.org). Three readings (replicates) were collected from each limb with a 1-minute interval between replicates.

Mechanical stimuli (EVF) were applied as ramped stimuli, with steadily increasing force applied until the behavioral response was elicited, or the maximum value was reached. A maximum value of 500g was set a priori prior to the study. Thresholds were measured in grams.

The hot stimuli were applied as fixed intensity stimuli. Hair over the site was clipped creating a 2 x 2 mm area prior to any testing. Hot stimuli represented probe settings of 46.5° C (cut-off time 20 s). The latency to respond was measured in seconds for thermal stimuli using a digital stopwatch (Seiko W073, Seiko, Japan) with the precision of one hundredth of a second.

### Activity monitors

Activity of the cats was measured on a subset of cats through the use of Actical activity monitors (Philips Respironics, Murrysville, PA) worn on the collar. The epoch was set at 1minute. Data were collected over a period of one week and levels of total activity and activity patterns compared between FIV infected and sham cats.

### Statistics

The design was optimized to evaluate effects of LM11A-31 on disease progression relative to placebo treated cats. This design was intended to match the prospective design of a trial of the compound in HIV infected humans. Disease progression from the 12-14 month test interval to the 18-20 month test interval was compared for FIV infected cats versus placebo controls. Cats were treated orally with LM11A-31, 13 mg/kg twice a day, (morning and evening approximately 12 hours apart), from 17.5 months to 20 months. The primary comparison across all procedures was drug vs placebo by t-test. Sham infected cats were used to provide an index of performance in uninfected cats as a secondary comparison to evaluate the extent of protection. If the data were not normally distributed, nonparametric analysis was applied. In three cases, cats were not included in the analysis as they failed to perform a task reliably, showed a behavior that was greater than 3 standard deviations from the mean of the other cats in the same group, or were not tested due to a medical condition that interfered with testing (such as gastrointestinal discomfort associated with hairballs and a paw injury). Specific comparisons are provided for each analysis.

## Acknowledgements

The authors would like to acknowledge the generous gift of LM11A-31 from PharmatrophiX which made this study possible.

